# Caffeine downregulates inflammatory pathways involved in autoimmunity

**DOI:** 10.1101/241539

**Authors:** Merve Iris, Pei-Suen Tsou, Amr H. Sawalha

**Author notes:** Please address correspondence to Amr H. Sawalha, MD; 5520 MSRB-1, SPC 5680, 1150 W. Medical Center Drive, Ann Arbor, MI 48109, USA.

## Abstract

**Objectives:** Caffeine is a widely consumed pharmacologically active product. In the present study, we focused on characterizing immunomodulatory effects of caffeine on peripheral blood mononuclear cells (PMBCs).

**Methods:** The effect of caffeine on gene expression profiles was initially evaluated using RNA sequencing data. Validation experiments were performed to confirm the results and examine dose-dependent effects of caffeine on PBMCs from healthy subjects. Gene expression levels were measured by real-time quantitative PCR, and cytokine production was determined using a multiplex cytokine assay.

**Results:** Caffeine at high doses showed a robust downregulatory effect of immune-related genes in PBMCs. Functional annotation analysis of downregulated genes revealed significant enrichment in cytokine activity and in genes related to several autoimmune diseases including lupus and rheumatoid arthritis. Dose-dependent validation experiments showed significant downregulation at the mRNA levels of key inflammatory genes including STAT1, TNF, and PPARG. TNF and PPARG were suppressed even with the lowest caffeine dose tested, which corresponds to the serum concentration of caffeine after administration of one cup of coffee. Cytokine levels of IL-8, MIP-1β, IL-6, IFN-γ, GM-CSF, TNF, IL-2, IL-4, MCP-1, and IL-10 were decreased significantly with caffeine treatment.

**Conclusion:** Our findings indicate potential downregulatory effects of caffeine on key inflammatory genes and cytokines, which play important role in autoimmunity. Further studies exploring therapeutic or disease-modulating potential of caffeine in autoimmune diseases and exploring the mechanisms involved are warranted.

## INTRODUCTION

Systemic autoimmune diseases result from dysregulation of the immune response which usually causes antigen-presenting cells to trigger autoreactive lymphocytes, and are often associated with chronic inflammation involving numerous cytokines. Persistent T cell activation and the production of inflammatory cytokines lead to tissue damage. The etiopathogenesis of autoimmune diseases is not completely understood. Many different combinations of genetic and environmental factors are thought to promote the development and progression of autoimmunity. The inheritance of epigenetic susceptibility loci can cause diversity in disease prevalence and severity between different ethnic and geographical populations [1, 2].

Previous studies have shown that epigenetic factors, including DNA methylation changes, are involved in the pathogenesis of autoimmune diseases including rheumatoid arthritis and systemic lupus erythematosus [3]. Being a source of external factors responsible for some epigenetic changes, diet has been shown to affect the severity of autoimmune diseases in animal models [4]. Moreover, the study of gene-environment interactions might help clarify the pathogenic consequences of both genetic and environmental influences in autoimmunity.

Caffeine is one of the most widely consumed pharmacologically active products in the world. Hence, its pharmacological properties, physiological effects, and potential treatment doses have been a subject of interest for years. Even though chocolates and cocoa containing foods store some amount of caffeine, the main source of caffeine in daily life is coffee or other caffeinated beverages such as energy drinks, tea, and carbonated soft drinks [5]. Caffeine has several effects on the body, including central nervous system stimulation. In addition, caffeine is a member of the methylxanthine family and has been used in neonatal intensive care units to treat apnea of prematurity. It has been studied for its effects on different diseases and was recently shown to reduce the incidence of bronchopulmonary dysplasia in neonates [6]. Previous studies suggested that caffeine is associated with reduced risk for some cancer types, chronic liver disease, Parkinson’s disease, Alzheimer’s disease, and diabetes [7, 8].

Studies using animal models suggest that caffeine has modulatory effects on elevated serum aminotransferase enzymes and the migration of inflammatory cells [8]. It also decreases serum inflammatory cytokine levels by suppressing their mRNA expression levels, suggesting an anti-inflammatory effect of caffeine [7, 9]. When studied with rats, chronic treatment with caffeine (10 and 30 mg/kg) has been found to ameliorate experimental autoimmune encephalomyelitis [10]. Methylxanthines, which include pentoxifylline and caffeine as members, mediate their effects by inhibiting cAMP-specific phosphodiesterase activity, thus, increasing levels of intracellular cAMP which has immunomodulatory effects on lymphocytes and monocytes/macrophages [11]. Cross-sectional studies have reported an association between higher coffee consumption and lower plasma concentrations of several markers of inflammation and endothelial dysfunction [12].

In this study, we focused on examining the immunomodulatory effects of caffeine using PBMCs obtained from healthy individuals. Our purpose was to evaluate the effect of caffeine on genes and pathways related to autoimmunity.

## RESULTS

To characterize the effect of caffeine on gene expression patterns in human PBMCs, we first used RNA sequencing data publically available from a recent study that examined gene expression profiles in multiple cell types, including primary human PBMCs, in response to a large number of environmental factors for the purpose of understanding gene-environment interactions [13]. Gene expression profiles in PBMCs from 3 healthy individuals stimulated with PHA (2.5ug/ml) with and without caffeine treatment (1.16 mM for 6 hours) were extracted. Differential expression was determined using a fold difference ≥1.5 and a p value ≤0.05 after correction for multiple testing using false discovery rate. We identified 3,437 upregulated and 2,811 downregulated genes with caffeine treatment (**Supplementary Tables 1 and 2**).

Gene ontology and pathway analysis revealed far more significant functional enrichment categories in genes downregulated compared to genes upregulated with caffeine treatment. Downregulated genes are clearly enriched in immune-related categories such as “cytokine activity” (GO:0005125) and “immune response” (GO:0006955). Likewise, when we examined disease enrichment patterns in gene downregulated with caffeine treatment we noticed downregulation of genes involved in multiple autoimmune diseases including systemic lupus erythematosus and rheumatoid arthritis with caffeine treatment (**Table 1**). Gene ontology analysis in upregulated genes pointed out “phosphotransferase activity” (GO:0016773) and “kinase activity” (GO:0016301), among other enriched gene ontologies (**Table 2**).

**Table 1:**
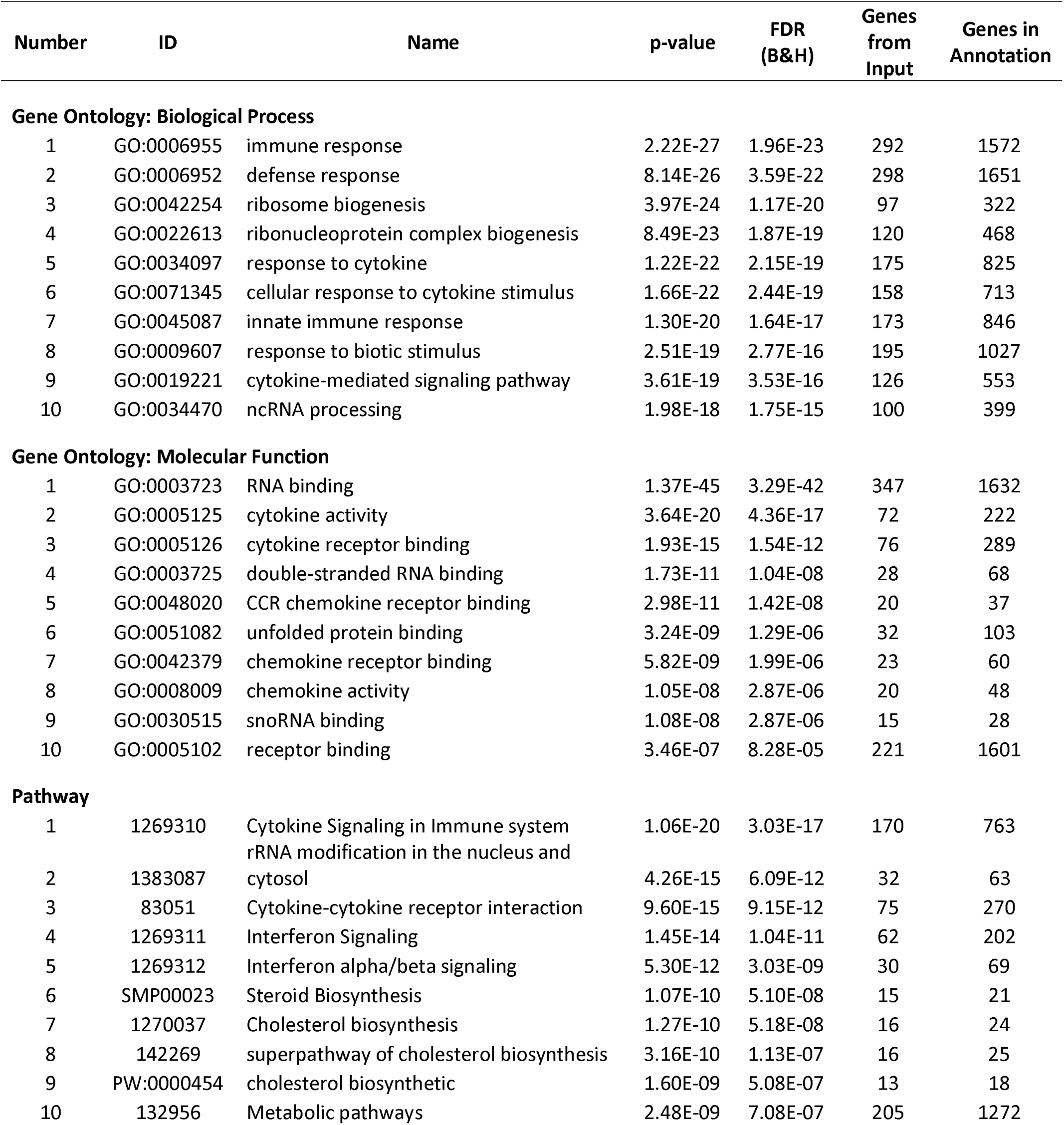
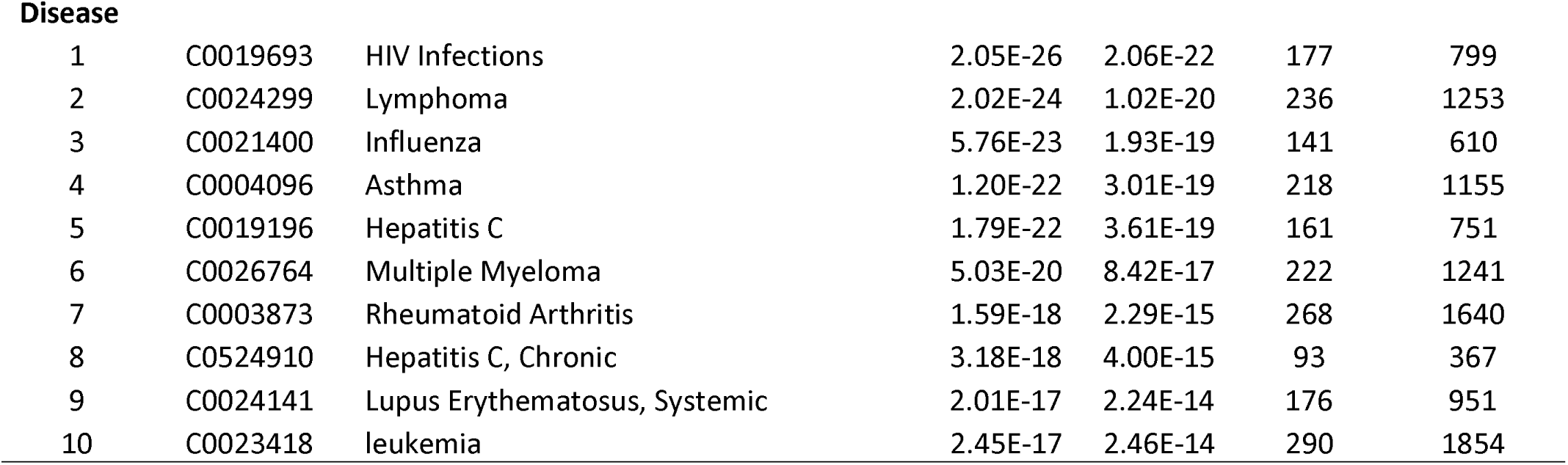
Gene Ontology-Biological Process, Gene Ontology-Molecular Function, Pathway, and Disease enrichment analysis in genes downregulated in PBMCs with caffeine treatment *in vitro*. Only the top 10 annotations in each category are shown.

| Number | ID | Name | p-value | FDR (B&H) | Genes from Input | Genes in Annotation |
| --- | --- | --- | --- | --- | --- | --- |
| **Gene Ontology: Biological Process** |  |  |  |  |  |  |
| 1 | GO:0006955 | immune response | 2.22E-27 | 1.96E-23 | 292 | 1572 |
| 2 | GO:0006952 | defense response | 8.14E-26 | 3.59E-22 | 298 | 1651 |
| 3 | GO:0042254 | ribosome biogenesis | 3.97E-24 | 1.17E-20 | 97 | 322 |
| 4 | GO:0022613 | ribonucleoprotein complex biogenesis | 8.49E-23 | 1.87E-19 | 120 | 468 |
| 5 | GO:0034097 | response to cytokine | 1.22E-22 | 2.15E-19 | 175 | 825 |
| 6 | GO:0071345 | cellular response to cytokine stimulus | 1.66E-22 | 2.44E-19 | 158 | 713 |
| 7 | GO:0045087 | innate immune response | 1.30E-20 | 1.64E-17 | 173 | 846 |
| 8 | GO:0009607 | response to biotic stimulus | 2.51E-19 | 2.77E-16 | 195 | 1027 |
| 9 | GO:0019221 | cytokine-mediated signaling pathway | 3.61E-19 | 3.53E-16 | 126 | 553 |
| 10 | GO:0034470 | ncRNA processing | 1.98E-18 | 1.75E-15 | 100 | 399 |
| **Gene Ontology: Molecular Function** |  |  |  |  |  |  |
| 1 | GO:0003723 | RNA binding | 1.37E-45 | 3.29E-42 | 347 | 1632 |
| 2 | GO:0005125 | cytokine activity | 3.64E-20 | 4.36E-17 | 72 | 222 |
| 3 | GO:0005126 | cytokine receptor binding | 1.93E-15 | 1.54E-12 | 76 | 289 |
| 4 | GO:0003725 | double-stranded RNA binding | 1.73E-11 | 1.04E-08 | 28 | 68 |
| 5 | GO:0048020 | CCR chemokine receptor binding | 2.98E-11 | 1.42E-08 | 20 | 37 |
| 6 | GO:0051082 | unfolded protein binding | 3.24E-09 | 1.29E-06 | 32 | 103 |
| 7 | GO:0042379 | chemokine receptor binding | 5.82E-09 | 1.99E-06 | 23 | 60 |
| 8 | GO:0008009 | chemokine activity | 1.05E-08 | 2.87E-06 | 20 | 48 |
| 9 | GO:0030515 | snoRNA binding | 1.08E-08 | 2.87E-06 | 15 | 28 |
| 10 | GO:0005102 | receptor binding | 3.46E-07 | 8.28E-05 | 221 | 1601 |
| **Pathway** |  |  |  |  |  |  |
| 1 | 1269310 | Cytokine Signaling in Immune system | 1.06E-20 | 3.03E-17 | 170 | 763 |
| 2 | 1383087 | rRNA modification in the nucleus and cytosol | 4.26E-15 | 6.09E-12 | 32 | 63 |
| 3 | 83051 | Cytokine-cytokine receptor interaction | 9.60E-15 | 9.15E-12 | 75 | 270 |
| 4 | 1269311 | Interferon Signaling | 1.45E-14 | 1.04E-11 | 62 | 202 |
| 5 | 1269312 | Interferon alpha/beta signaling | 5.30E-12 | 3.03E-09 | 30 | 69 |
| 6 | SMP00023 | Steroid Biosynthesis | 1.07E-10 | 5.10E-08 | 15 | 21 |
| 7 | 1270037 | Cholesterol biosynthesis | 1.27E-10 | 5.18E-08 | 16 | 24 |
| 8 | 142269 | superpathway of cholesterol biosynthesis | 3.16E-10 | 1.13E-07 | 16 | 25 |
| 9 | PW:0000454 | cholesterol biosynthetic | 1.60E-09 | 5.08E-07 | 13 | 18 |
| 10 | 132956 | Metabolic pathways | 2.48E-09 | 7.08E-07 | 205 | 1272 |
| **Disease** |  |  |  |  |  |  |
| 1 | C0019693 | HIV Infections | 2.05E-26 | 2.06E-22 | 177 | 799 |
| 2 | C0024299 | Lymphoma | 2.02E-24 | 1.02E-20 | 236 | 1253 |
| 3 | C0021400 | Influenza | 5.76E-23 | 1.93E-19 | 141 | 610 |
| 4 | C0004096 | Asthma | 1.20E-22 | 3.01E-19 | 218 | 1155 |
| 5 | C0019196 | Hepatitis C | 1.79E-22 | 3.61E-19 | 161 | 751 |
| 6 | C0026764 | Multiple Myeloma | 5.03E-20 | 8.42E-17 | 222 | 1241 |
| 7 | C0003873 | Rheumatoid Arthritis | 1.59E-18 | 2.29E-15 | 268 | 1640 |
| 8 | C0524910 | Hepatitis C, Chronic | 3.18E-18 | 4.00E-15 | 93 | 367 |
| 9 | C0024141 | Lupus Erythematosus, Systemic | 2.01E-17 | 2.24E-14 | 176 | 951 |
| 10 | C0023418 | leukemia | 2.45E-17 | 2.46E-14 | 290 | 1854 |

**Table 2:**
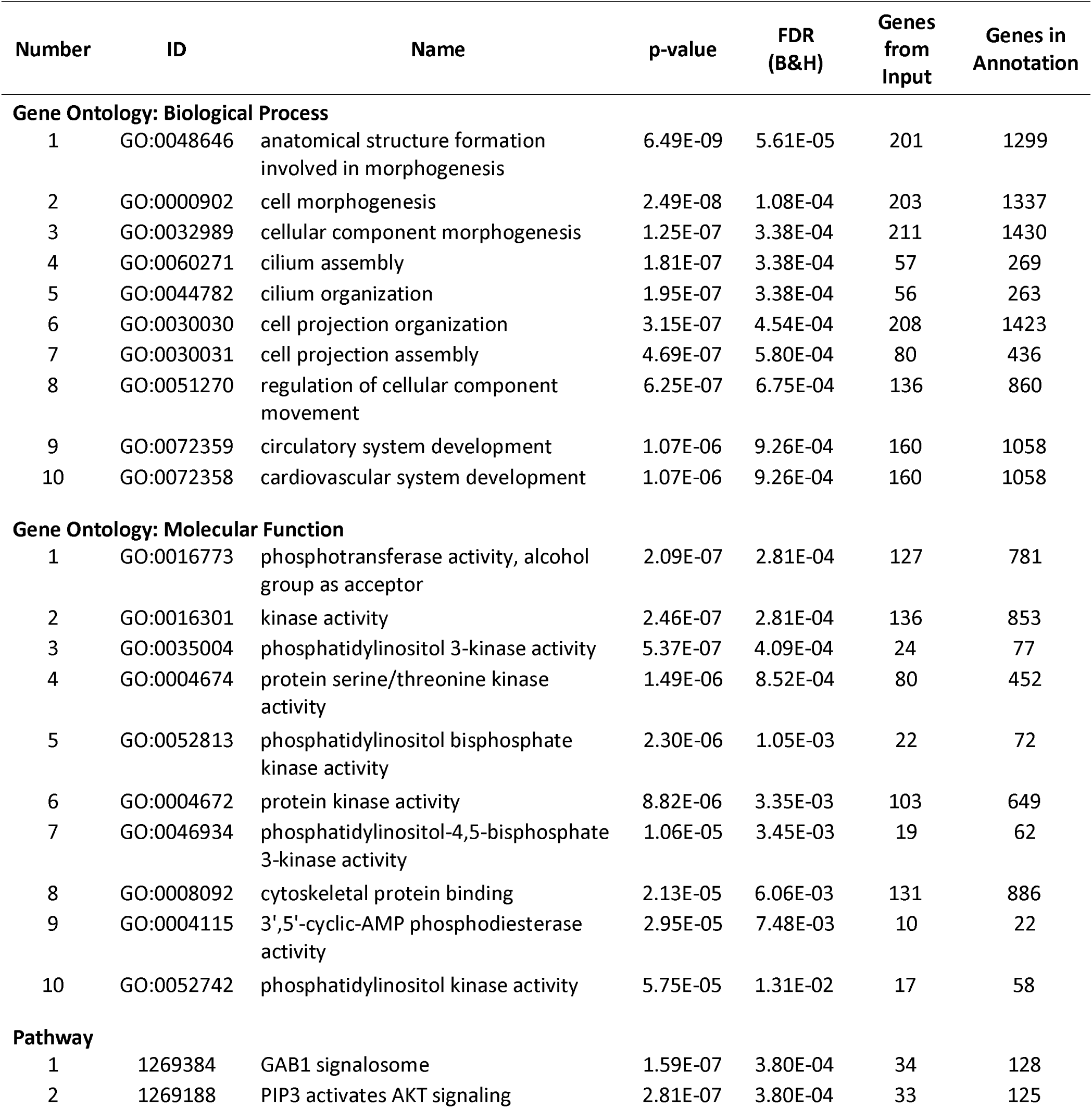
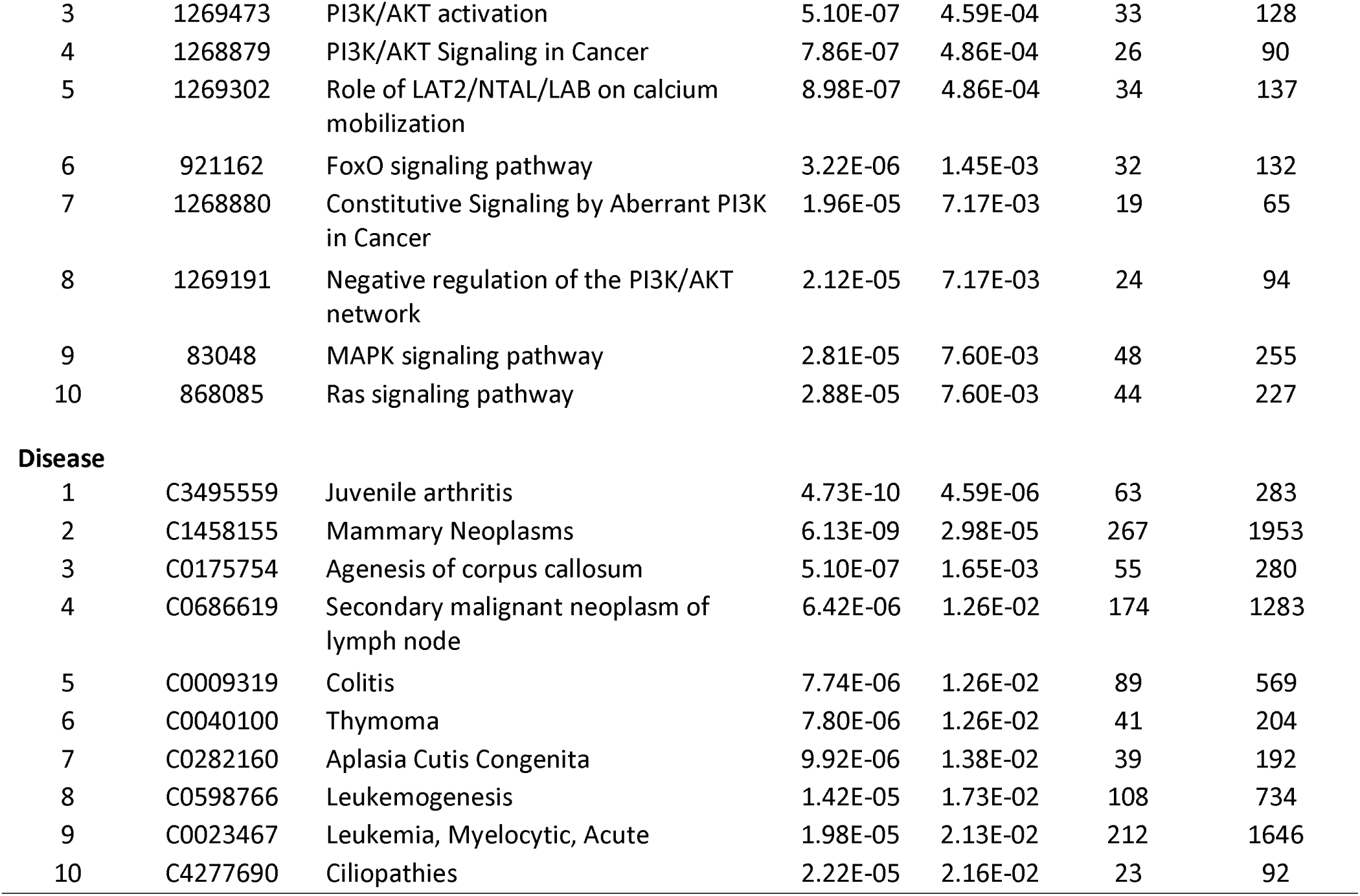
Gene Ontology-Biological Process, Gene Ontology-Molecular Function, Pathway, and Disease enrichment analysis in genes upregulated in PBMCs with caffeine treatment *in vitro*. Only the top 10 annotations in each category are shown.

| Number | ID | Name | p-value | FDR (B&H) | Genes from Input | Genes in Annotation |
| --- | --- | --- | --- | --- | --- | --- |
| **Gene Ontology: Biological Process** |  |  |  |  |  |  |
| 1 | GO:0048646 | anatomical structure formation involved in morphogenesis | 6.49E-09 | 5.61E-05 | 201 | 1299 |
| 2 | GO:0000902 | cell morphogenesis | 2.49E-08 | 1.08E-04 | 203 | 1337 |
| 3 | GO:0032989 | cellular component morphogenesis | 1.25E-07 | 3.38E-04 | 211 | 1430 |
| 4 | GO:0060271 | cilium assembly | 1.81E-07 | 3.38E-04 | 57 | 269 |
| 5 | GO:0044782 | cilium organization | 1.95E-07 | 3.38E-04 | 56 | 263 |
| 6 | GO:0030030 | cell projection organization | 3.15E-07 | 4.54E-04 | 208 | 1423 |
| 7 | GO:0030031 | cell projection assembly | 4.69E-07 | 5.80E-04 | 80 | 436 |
| 8 | GO:0051270 | regulation of cellular component movement | 6.25E-07 | 6.75E-04 | 136 | 860 |
| 9 | GO:0072359 | circulatory system development | 1.07E-06 | 9.26E-04 | 160 | 1058 |
| 10 | GO:0072358 | cardiovascular system development | 1.07E-06 | 9.26E-04 | 160 | 1058 |
| **Gene Ontology: Molecular Function** |  |  |  |  |  |  |
| 1 | GO:0016773 | phosphotransferase activity, alcohol group as acceptor | 2.09E-07 | 2.81E-04 | 127 | 781 |
| 2 | GO:0016301 | kinase activity | 2.46E-07 | 2.81E-04 | 136 | 853 |
| 3 | GO:0035004 | phosphatidylinositol 3-kinase activity | 5.37E-07 | 4.09E-04 | 24 | 77 |
| 4 | GO:0004674 | protein serine/threonine kinase activity | 1.49E-06 | 8.52E-04 | 80 | 452 |
| 5 | GO:0052813 | phosphatidylinositol bisphosphate kinase activity | 2.30E-06 | 1.05E-03 | 22 | 72 |
| 6 | GO:0004672 | protein kinase activity | 8.82E-06 | 3.35E-03 | 103 | 649 |
| 7 | GO:0046934 | phosphatidylinositol-4,5-bisphosphate 3-kinase activity | 1.06E-05 | 3.45E-03 | 19 | 62 |
| 8 | GO:0008092 | cytoskeletal protein binding | 2.13E-05 | 6.06E-03 | 131 | 886 |
| 9 | GO:0004115 | 3',5'-cyclic-AMP phosphodiesterase activity | 2.95E-05 | 7.48E-03 | 10 | 22 |
| 10 | GO:0052742 | phosphatidylinositol kinase activity | 5.75E-05 | 1.31E-02 | 17 | 58 |
| **Pathway** |  |  |  |  |  |  |
| 1 | 1269384 | GAB1 signalosome | 1.59E-07 | 3.80E-04 | 34 | 128 |
| 2 | 1269188 | PIP3 activates AKT signaling | 2.81E-07 | 3.80E-04 | 33 | 125 |
| 3 | 1269473 | PI3K/AKT activation | 5.10E-07 | 4.59E-04 | 33 | 128 |
| 4 | 1268879 | PI3K/AKT Signaling in Cancer | 7.86E-07 | 4.86E-04 | 26 | 90 |
| 5 | 1269302 | Role of LAT2/NTAL/LAB on calcium mobilization | 8.98E-07 | 4.86E-04 | 34 | 137 |
| 6 | 921162 | FoxO signaling pathway | 3.22E-06 | 1.45E-03 | 32 | 132 |
| 7 | 1268880 | Constitutive Signaling by Aberrant PI3K in Cancer | 1.96E-05 | 7.17E-03 | 19 | 65 |
| 8 | 1269191 | Negative regulation of the PI3K/AKT network | 2.12E-05 | 7.17E-03 | 24 | 94 |
| 9 | 83048 | MAPK signaling pathway | 2.81E-05 | 7.60E-03 | 48 | 255 |
| 10 | 868085 | Ras signaling pathway | 2.88E-05 | 7.60E-03 | 44 | 227 |
| **Disease** |  |  |  |  |  |  |
| 1 | C3495559 | Juvenile arthritis | 4.73E-10 | 4.59E-06 | 63 | 283 |
| 2 | C1458155 | Mammary Neoplasms | 6.13E-09 | 2.98E-05 | 267 | 1953 |
| 3 | C0175754 | Agenesis of corpus callosum | 5.10E-07 | 1.65E-03 | 55 | 280 |
| 4 | C0686619 | Secondary malignant neoplasm of lymph node | 6.42E-06 | 1.26E-02 | 174 | 1283 |
| 5 | C0009319 | Colitis | 7.74E-06 | 1.26E-02 | 89 | 569 |
| 6 | C0040100 | Thymoma | 7.80E-06 | 1.26E-02 | 41 | 204 |
| 7 | C0282160 | Aplasia Cutis Congenita | 9.92E-06 | 1.38E-02 | 39 | 192 |
| 8 | C0598766 | Leukemogenesis | 1.42E-05 | 1.73E-02 | 108 | 734 |
| 9 | C0023467 | Leukemia, Myelocytic, Acute | 1.98E-05 | 2.13E-02 | 212 | 1646 |
| 10 | C4277690 | Ciliopathies | 2.22E-05 | 2.16E-02 | 23 | 92 |

As we detected a strong evidence for downregulating pathways involved in many immune-related genes with caffeine treatment, we performed validation experiments in a selected number of relevant downregulated genes to examine the dose-dependent effect of caffeine concentrations. We focused on the potential downregulatory effects of caffeine on autoimmunity and related inflammatory pathways. We selected genes within the most downregulated functional categories including “Cytokine Activity” (GO:0005125), “Immune Response” (GO:0006955), “Cytokine Signaling in Immune System” (ID:1269310), “Cytokine-cytokine Receptor Interaction” (ID: 83051) and “Lupus Erythematosus, Systemic” (ID: C0024141). We investigated the mRNA expression of MX1, STAT1, IRF5, IFNG, PPARG and TNF in PBMCs isolated from healthy normal blood donors, with and without low, intermediate, and high caffeine concentrations. These concentrations correspond to the mean value of Cmax in serum when a person drinks one cup of coffee, the maximum therapeutic dose of caffeine, and the dose used in the previous RNA sequencing experiment for the purpose of understanding gene-environment interactions, respectively.

We observed significant downregulation of STAT1, TNF, and PPARG with caffeine treatment in PBMCs (**Table 3**). A dose-dependent reduction in the expression STAT1 was observed with intermediate (p=0.0384) and high (p=0.0001) caffeine concentrations. Expression levels of TNF and PPARG were significantly reduced at all caffeine concentrations, with a dose-response effect also observed (**Figure 1**). In contrast to STAT1, mRNA expression levels of TNF and PPARG were also suppressed at low caffeine concentration (0.019 mM), which is approximately equivalent to the serum caffeine concentration after administration of one cup of coffee.

**Table 3:** Quantitative real time RT PCR results in genes downregulated with caffeine by RNA sequencing data and selected for confirmation using different caffeine concentrations. The fold difference ± standard deviation is listed relative to untreated cells and the P values were adjusted using Dunnett’s multiple comparisons test.

| **Gene** | **Expression levels with caffeine treatment compared to untreated cells** |  |  |  |  |  |
| --- | --- | --- | --- | --- | --- | --- |
|  | **Low** |  | **Medium** |  | **High** |  |
|  | Fold change | Adjusted p value | Fold change | Adjusted p value | Fold change | Adjusted p value |
| *TNF* | 0.731±0.326 | **0.023** | 0.625±0.302 | **0.0015** | 0.279±0.042 | **0.0001** |
| *PPARG* | 0.605±0.377 | **0.011** | 0.474±0.303 | **0.0007** | 0.415±0.283 | **0.0002** |
| *STAT1* | 0.885±0.175 | 0.15 | 0.845±0.112 | **0.038** | 0.527±0.193 | **0.0001** |
| *MX1* | 1.151±0.238 | 0.54 | 1.033±0.249 | 0.99 | 1.336±0.524 | **0.043** |
| *IRF5* | 0.769±0.24 | 0.14 | 0.896±0.34 | 0.7 | 0.949±0.343 | 0.94 |
| *IFNG* | 0.944±0.256 | 0.9 | 0.898±0.328 | 0.63 | 0.651±0.24 | **0.0054** |

**Figure 1:**
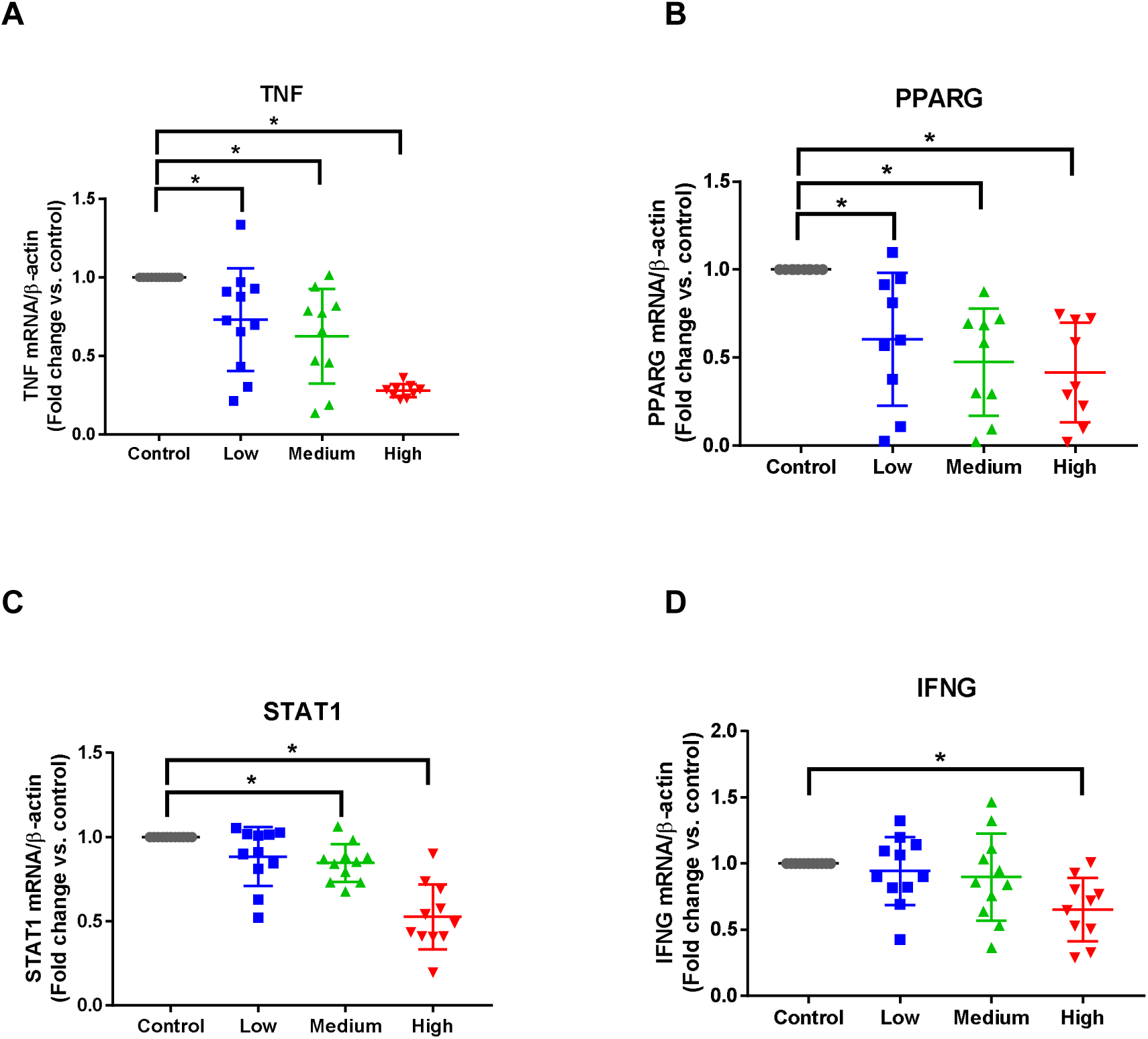
Quantitative real time RT-PCR results for TNF, PPARG, and STAT1 in PBMCs with and without caffeine treatment *in vitro* demonstrates a dose-dependent downregulation in gene expression level. **A)** TNF mRNA expression (p=0.023, p=0.0015, and p=0.0001 for low, medium, and high caffeine concentrations compared to untreated cells, respectively) (One-way ANOVA p<0.0001). **B)** PPARG mRNA expression (p=0.0105, p=0.0007, and p=0.0002 for low, medium, and high caffeine concentrations compared to untreated cells, respectively) (One-way ANOVA p=0.0002). **C)** STAT1 mRNA expression p=0.15, p=0.0384, and p=0.0001 for low, medium, and high caffeine concentrations compared to untreated cells, respectively) (One-way ANOVA p<0.0001). **D)** IFNG mRNA expression p=0.9, p=0.63, and p=0.0054 for low, medium, and high caffeine concentrations compared to untreated cells, respectively) (One-way ANOVA p=0.011).

Because of enrichment of cytokine genes and cytokine activity in functional enrichment analysis of genes downregulated with caffeine treatment, we measured cytokine production levels in cell culture media from PBMCs treated with and without caffeine. We observed significant reduction in the production of multiple inflammatory cytokines, validating gene expression patterns we observed. IL-8 levels were significantly reduced with any caffeine treatment, in a dose-dependent manner. Significant reduction in the levels of IL-6, IFNγ, TNF, MCP-1 and IL-10 was also observed at medium and high caffeine concentrations (**Figure 2**, **Table 4**).

**Figure 2:**
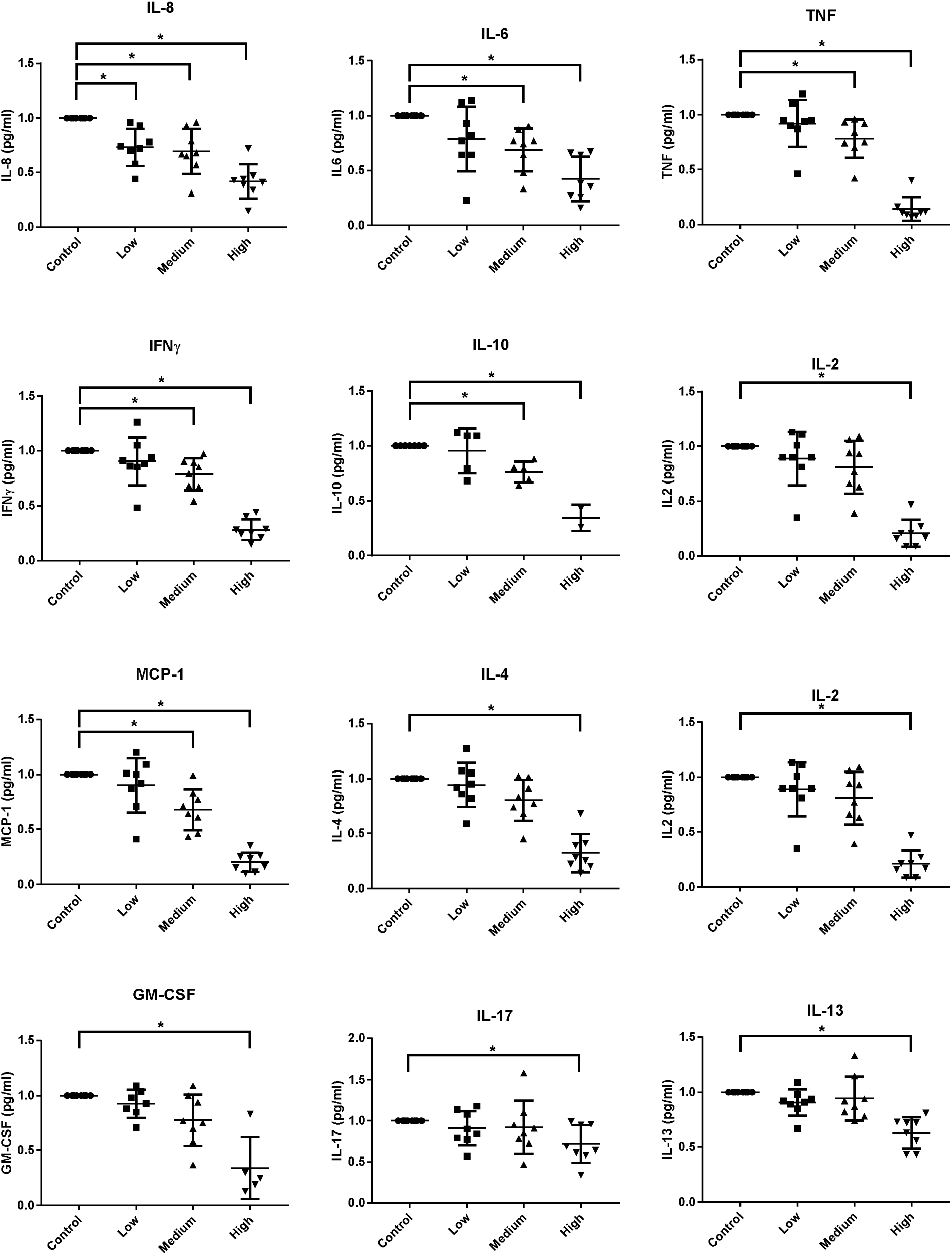
Cytokine levels measured in the supernatants of PBMCs with and without caffeine treatment. *, p-value <0.05.

**Table 4:**
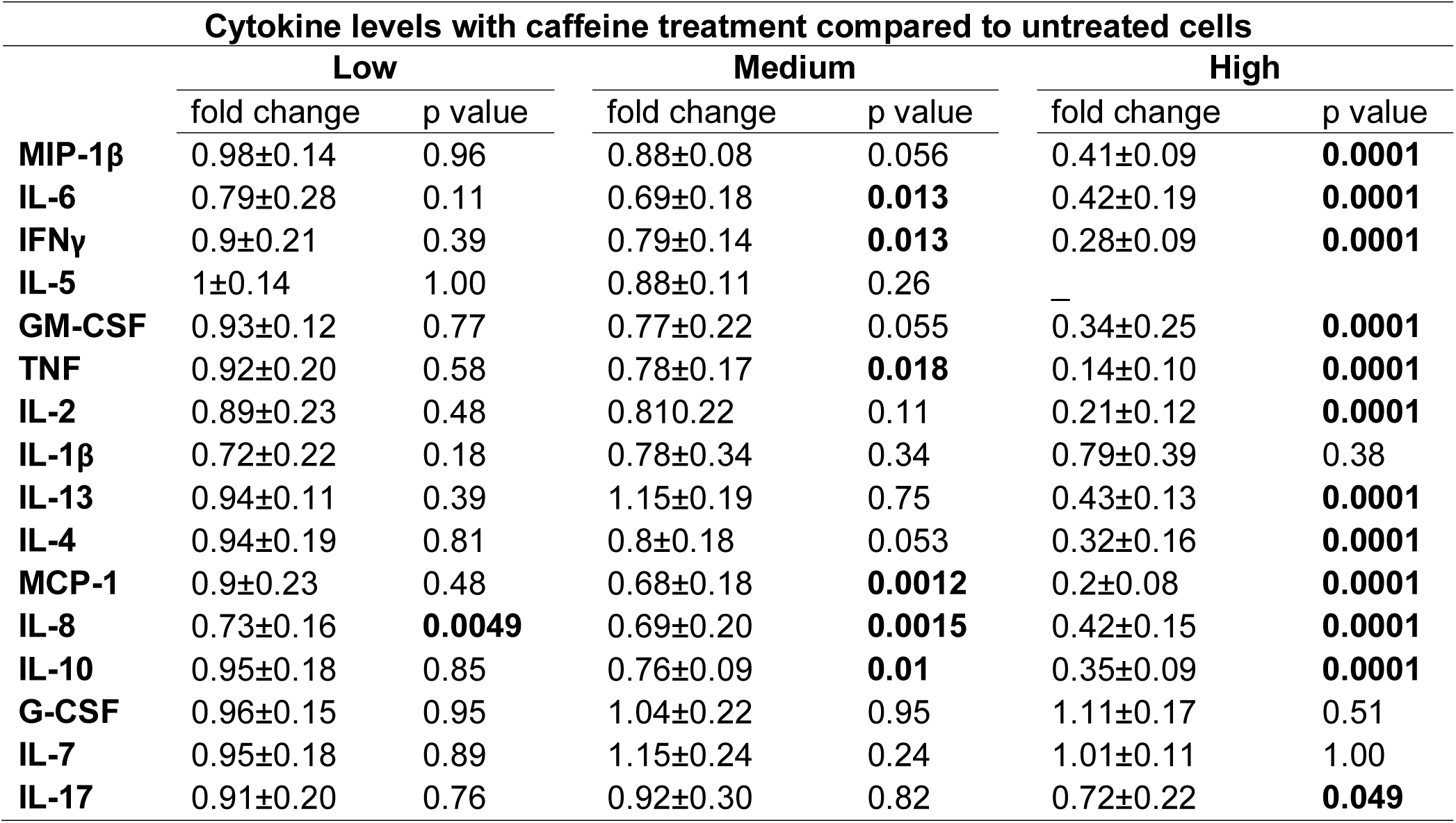
Cytokine levels measured in the supernatants of PBMCs, using different caffeine concentrations. The fold difference ± standard deviation is listed relative to untreated cells and the P values were adjusted using Dunnett’s multiple comparisons test.

## DISCUSSION

Many autoimmune diseases accompany chronic inflammatory state which involves multiple organs and systems. According to National Institute of Allergy and Infectious Diseases, there are more than 80 autoimmune diseases ranging from mild to severe multiorgan involving forms and affecting patients’ quality of life and mental status [14]. In severe forms, they can be related to increased mortality by causing end-organ failure.

Heritable genetic alterations, immune system abnormalities, and environmental elements contribute to the development and progression of many autoimmune diseases [15]. Indeed, environmental factors can influence disease susceptibility by means of epigenetics mechanisms such as DNA methylation and histone modification processes. In this regard, it is not unexpected that diet might have an important role. Recent studies have demonstrated the influence of dietary factors on autoantibody levels and disease severity in mouse models of autoimmunity [4, 16].

The present study questioned and demonstrated the *in vitro* downregulatory effects of a commonly consumed product, caffeine, on inflammation and autoimmunity related genes, pathways, and inflammatory cytokine levels.

TNF, being an important inflammatory cytokine in many immune-mediated diseases, is target of several therapeutics [17]. Previous studies indicated that TNF is involved in the release of other inflammatory cytokines such as IL-1 and IL-6 [18]. We demonstrate suppression of TNF production both at the mRNA and protein levels with low to intermediate caffeine concentrations. Indeed, caffeine concentrations corresponding to serum levels after the consumption of one cup of coffee was associated with significant suppression of TNF production in PBMCs.

Cytokines are known to have a key role in autoimmune diseases, and several pro-inflammatory cytokines were reduced with caffeine treatment in our study. IL-6 and IL-10 which are both significantly elevated in the serum with increased disease activity in lupus patients can be downregulated with caffeine. Indeed, IL-10 is produced in increased amounts by lupus PBMCs and IL-10 levels have been shown to correlated with disease activity in lupus patients [19].

Previous studies have demonstrated higher levels of IFNγ mRNA in peripheral blood T cells from patients with lupus compared to normal controls after CD28 costimulation [20]. Moreover, in lupus monocytes IFNγ signaling has been found to be more effective in inducing STAT1 phosphorylation compared with monocytes from healthy individuals [21]. It has been also reported that lupus patients have elevated STAT1 and IFNγ expression in PBMCs and that this elevation correlates with disease activity [20, 21]. Cytokines that signal through STATs are crucial in host defense and inflammation, and overactivation of STATs can induce immune pathologies [22]. Our data demonstrate significant down regulation of both STAT1 and IFNγ with caffeine.

In summary, we demonstrate that caffeine, even with low concentrations, can suppress gene expression of pro-inflammatory genes and cytokines that play key roles in autoimmune diseases. Further work to characterize the mechanisms involved in the anti-inflammatory effects observed and whether caffeine supplementation might have beneficial effects in patients with autoimmune diseases are worthy of investigation.

## METHODS

### RNA sequencing data

RNA sequencing data derived from PMBCs with and without treatment with caffeine were extracted from a publically available dataset. These data were generated using PBMCs isolated from 3 healthy individuals. PBMCs were activated with PHA and then treated with or without caffeine for 6 hours at a concentration of 1.16 mM prior to RNA expression and sequencing [13].

### Isolating PBMCs and cell culture

PBMCs were isolated using density gradient centrifugation (Ficoll) from buffy coats isolated from whole blood of 11 healthy blood donors and stored in liquid nitrogen. PBMCs were thawed and suspended in warm RPMI (RPMI+L-glutamine+10% FBS)/benzonase. Following resuspension, cells were centrifuged at 300xg for 10 minutes, suspended, then centrifuged again at 100xg for 10 minutes. PBMCs were then cultured in RPMI without benzonase and transferred to 6-well plates to rest overnight at 37 degrees Celsius with 5.0% CO_2_. Cells were then centrifuged at 300xg for 10 minutes, then plated at 1×10^6^ cells/mL in RPMI with L-Glutamine and 0.1% charcoal-stripped FBS (1–2 million cells in 1ml media in a 12-well plate). Cells were activated with PHA (2.5µg/mL) and incubated with or without caffeine for 6 hours. Three different concentration of caffeine were used: Low (0.019 mM), Intermediate (0.102 mM), and high (1.16 mM). The low dose corresponds to the mean value of Cmax in serum when a person drinks one cup of hot coffee containing 160 mg of caffeine [23]. For the intermediate dose, we used caffeine at the maximum therapeutic concentration according to Mayo Medical Laboratories (www.mayomedicallaboratories.com). The high dose was the dose used in a previous experiment for the purpose of understanding gene-environment interactions [13]. Only the highest dose was within what is considered toxic levels of serum caffeine concentration (Mayo Medical Laboratories).

### Preparation of RNA and qPCR analysis

Cells were centrifuged at 300xg for 10 minutes, and cell culture media were stored at −80 degrees Celsius for subsequent cytokine analysis. The cells were washed with PBS then TRIzol Reagent (Thermo Fisher Scientific) was added. RNA was extracted using Direct-zol RNA Isolation Kit (Zymo Research). cDNA was prepared using the Verso cDNA Synthesis Kit (Thermo Fisher Scientific). Quantitative reverse transcriptase PCR was performed using primers for human MX1, STAT1, IRF5, IFNG, PPARG, TNF and β-actin. Power SYBR Green PCR master mix (Applied Biosystems) was used for qPCR, which was run by a ViiA™ 7 Real-Time PCR System. Primers were obtained from KiCqStart^®^ SYBR^®^ Green Primers from Sigma and QuantiTect Primer Assays from Qiagen.

### Bioinformatics analysis

Functional enrichment analysis of genes upregulated and downregulated with caffeine treatment was performed using ToppGene to identify Gene Ontology (Molecular Function and Biological Process), Pathway, and Disease annotations [24].

### Cytokine analysis

Cytokines were measured using Bio-Plex Pro™ Human Cytokine 17-Plex Assay which includes MIP-1β, IL-6, IFNγ, IL-5, GM-CSF, TNF, IL-2, IL-1β, IL-13, IL-4, MCP-1, IL-8, IL-10, G-CSF, IL-7, IL-12, and IL-17.

### Statistical analysis

Statistical analysis was performed using GraphPad Prism 7.00 software. To compare the differences between groups, we used ANOVA with post-hoc test, with statistical significance at a P value <0.05.

